# Longitudinal development of hippocampal subregions from childhood to adulthood

**DOI:** 10.1101/186270

**Authors:** Christian K. Tamnes, Marieke G. N. Bos, Ferdi C. van de Kamp, Sabine Peters, Eveline A. Crone

**Author notes:** Corresponding author: Christian K. Tamnes, Department of Psychology, University of Oslo, PO Box 1094 Blindern, 0317 Oslo, Norway.

## Abstract

Detailed descriptions of the development of the hippocampus promise to shed light on the neural foundation of development of memory and other cognitive functions, as well as the emergence of major mental disorders. Hippocampus is a heterogeneous structure with a well characterized internal complexity, but development of its distinct subregions in humans has remained poorly described. We analyzed magnetic resonance imaging (MRI) data from a large longitudinal sample (270 participants, 678 scans) using an automated segmentation tool and mixed models to delineate the development of hippocampal subregion volumes from childhood to adulthood. We also examined sex differences in subregion volumes and their development, and associations between hippocampal subregions and general cognitive ability. Nonlinear developmental trajectories with early volume increases were observed for subiculum, cornu ammonis (CA) 1, molecular layer (ML) and fimbria. In contrast, parasubiculum, presubiculum, CA2/3, CA4 and the granule cell layer of the dentate gyrus (GC-DG) showed linear volume decreases. No sex differences were found in hippocampal subregion development. Finally, general cognitive ability was positively associated with CA2/3 and CA4 volumes, as well as with ML development. In conclusion, hippocampal subregions appear to develop in diversified ways across adolescence, and specific subregions may link to general cognitive level.

**Highlights:**

- Hippocampal subregions develop in differential ways from childhood to adulthood
- Subiculum, CA1, ML and fimbria showed nonlinear trajectories with initial increases
- Parasubiculum, presubiculum, CA2/3, CA4 and GC-DG showed linear volume decreases
- There were no sex differences in hippocampal subregion development
- General cognitive ability associated with CA2/3 and CA4 volumes and ML development

## Introduction

Knowledge of the development of the hippocampus from childhood to adulthood is important for understanding the neural foundation of development of cognitive functions, including episodic memory (Ghetti & Bunge, 2012; Østby, Tamnes, Fjell, & Walhovd, 2012). Moreover, it may offer insight into the origin and ontogeny of major mental disorders including schizophrenia and depression, which frequently emerge in adolescence (Lee, Heimer, et al., 2014; Whiteford et al., 2013), and for which the hippocampus appears to be a key node in the underlying distributed brain networks (Schmaal et al., 2016; van Erp et al., 2016). Magnetic resonance imaging (MRI) studies have investigated age-related differences or longitudinal changes in hippocampal volume in children and adolescents. The hippocampus is however not a uniform structure, but contains anatomically and functionally distinct regions (Amaral & Lavenex, 2007). It is thus possible that different subregions develop differently.

Hippocampal volume increases during childhood (Brown et al., 2012; Gilmore et al., 2012; Hu, Pruessner, Coupe, & Collins, 2013; Swagerman, Brouwer, de Geus, Hulshoff Pol, & Boomsma, 2014; Uematsu et al., 2012), but results for the adolescent period have been more variable. Several cross-sectional studies (Koolschijn & Crone, 2013; Muftuler et al., 2011; Yurgelun-Todd, Killgore, & Cintron, 2003; Østby et al., 2009) and some longitudinal studies (Mattai et al., 2011; Sullivan et al., 2011) found no significant age effects. More recent longitudinal studies have found volume increase (Dennison et al., 2013), decrease (Tamnes et al., 2013), or a quadratic inverted U-shaped trajectory (Narvacan, Treit, Camicioli, Martin, & Beaulieu, 2017; Wierenga et al., 2014). The latter finding is supported by a recent multisite longitudinal developmental study (Herting et al., 2018) and a large cross-sectional lifespan study (Coupe, Catheline, Lanuza, Manjon, & Alzheimer’s Disease Neuroimaging Initiative, 2017).

Estimating whole hippocampal volume may however mask regional developmental differences. Anatomically, the hippocampus is a unique structure consisting of cytoarchitectonically distinct subregions, including the cornu ammonis (CA) subfields, the dentate gyrus (DG) and the subicular complex (Insausti & Amaral, 2012). The hippocampal formation also has a unique set of largely unidirectional, excitatory pathways along the transverse plane (Amaral & Lavenex, 2007). Despite this well characterized internal complexity, researchers studying the human hippocampus *in vivo* have traditionally modelled and measured it as a whole (but see (Insausti, Cebada-Sanchez, & Marcos, 2010)). Novel protocols to segment the hippocampal subregions in MRI images have however been developed. Analysis of subregion within the hippocampus may unravel heterogeneous developmental patterns with differential functional relevance.

A pioneer study indicated different developmental changes in subareas of the hippocampus, mainly with increases in posterior areas and decreases in anterior areas (Gogtay et al., 2006). This was partly supported by a study investigating age-related differences in the head, body and tail of the hippocampus, finding an increase in the volume of the body and decreases in the right head and tail (DeMaster, Pathman, Lee, & Ghetti, 2014). Other studies have investigating the development of more clearly defined hippocampal subregions, including its subfields. Krogsrud et al. (2014) found that most subregions showed age-related volume increases from early childhood until approximately 13-15 years, followed by little differences. For a subsample of these participants, Tamnes et al. (2014) performed a longitudinal follow-up and found that change rates were different across subregions, but that nearly all showed small volume decreases in the teenage years. Combined, these results fit with the observed inverted U-shaped trajectory for whole hippocampal volume. Based on manual segmentation of subfields in the hippocampus body, Lee et al. (2014) found age-related increases in the right CA1 and CA3/DG volumes into early adolescence. Finally, in a lifespan sample, Daugherty et al. (2016) performed manual tracing on slices in the anterior hippocampus body and found negative relationships with age during development for CA1/2 and CA3/DG volumes.

Together, these results suggest that hippocampal subregions continue to change in subtle and diverse ways through childhood and adolescence, but the available studies have major limitations. First, several of the studies had relatively small samples. Second, only two of the studies had longitudinal data (Gogtay et al., 2006; Tamnes et al., 2014) and could investigate growth trajectories. Third, two of the previous studies (Krogsrud et al., 2014; Tamnes et al., 2014) used an automated segmentation procedure (Van Leemput et al., 2009) for which the reliability and validity has later been challenged (de Flores et al., 2015; Wisse, Biessels, & Geerlings, 2014), and these results have to be interpreted with caution. The other two studies of specific subregions (Daugherty et al., 2016; Lee, Ekstrom, et al., 2014) used manual tracing protocols (Ekstrom et al., 2009; Mueller et al., 2007) which yield estimates of a smaller number regions measured only in the hippocampal body. Moreover, manual segmentation is laborious and can be infeasible for large longitudinal studies, and also requires some subjectivity and is thus vulnerable to bias (Schlichting, Mack, Guarine, & Preston, 2017). The manual methods are thus not optimal in the context of the increasing focus on larger samples to obtain adequate statistical power (Button et al., 2013) and open science and reproducibility (Nichols et al., 2017). On the other hand, however, automated methods have potential limitations related to validity, e.g., the segmentation tool can be biased towards a different age group or a different type of sample (see limitations section).

We aimed to partially address some of the shortcomings of the previous studies by analyzing data from a large longitudinal sample of 270 participants with 678 MRI scans in the age-range 8-28 years using a novel automated segmentation tool. Specifically, we aimed to characterize the development of hippocampal subregion volumes from childhood to adulthood. Second, previous studies of sex differences in hippocampal development have been inconsistent (Herting et al., 2018), so we aimed to investigate whether hippocampal subregion volumes and development differs between girls and boys. Finally, we aimed to investigate how hippocampal subregions related to general cognitive ability, which previous studies have found to be related to cortical and white matter structure and development (Shaw et al., 2006; Tamnes et al., 2010; Walhovd et al., 2016).

## Materials and Methods

### Procedure and participants

The current study was part of the accelerated longitudinal research project *Braintime* (Becht et al., in press; Bos, Peters, van de Kamp, Crone, & Tamnes, in press; Peters & Crone, 2017; Schreuders et al., in press) performed in Leiden, the Netherlands, and approved by the Institutional Review Board at Leiden University Medical Center. Hippocampal subregions have not previously been analyzed in this project. At each time-point (TP), informed consent was obtained from each participant or from a parent in case of minors. Participants received presents and parents received financial reimbursement for travel costs. The participants were recruited through local schools and advertisements across Leiden, The Netherlands. All included participants were required to be fluent in Dutch, right-handed, have normal or corrected-to-normal vision, and to not report neurological or mental health problems or use of psychotropic medication. An initial sample of 299 participants (153 females, 146 males) in the age range 8-26 years old was recruited. All participants were invited to participate in three consecutive waves of data collection approximately two years apart. General cognitive ability was estimated at TP1 and TP2 using different subtests from age-appropriate Wechsler Intelligence Scales (WISC and WAIS) to avoid practice effects; TP1: Similarities and Block Design; TP2: Picture Completion and Vocabulary; TP3: no measurement. All included participants had an estimated IQ ≥ 80.

The final sample for the current study consisted of participants who had at least one structural MRI scan that was successfully processed through both the standard and hippocampal subfield segmentation longitudinal pipelines of FreeSurfer and which passed our quality control (QC) procedure (see below). This yielded a dataset consisting of 270 participants (145, females, 125 males) with 678 scans (Table 1); 169 participants had scans from 3 TPs, 70 participants had scans from two TPs, and 31 participants had one scan. The mean number of scans per participants was 2.51 (SD = 0.69). The mean interval for longitudinal follow-up scans in the final dataset was 2.11 years (SD = 0.46, range = 1.55-4.43).

**Table 1.** Sample characteristics for each time-point (TP)

|  | TP1 | TP2 | TP3 |
| --- | --- | --- | --- |
| n | 237 | 224 | 217 |
| n females/males | 128/109 | 118/106 | 119/98 |
| Age, mean (SD) | 14.5 (3.7) | 16.4 (3.6) | 18.4 (3.7) |
| Age, range | 8.0 – 26.0 | 9.9 – 26.6 | 11.9 – 28.7 |
| Estimated IQ, mean (SD) | 110.0 (10.2) | 108.5 (10.1)<sup>a</sup> | – |
| Estimated IQ, range | 80 – 138 | 80 – 148<sup>a</sup> | – |

<sup>a</sup> data missing for 1 participant

### Image acquisition

All scanning was performed on a single 3-Tesla Philips Achieve whole body scanner, using a 6 element SENSE receiver head coil (Philips, Best, The Netherlands) at Leiden University Medical Centre. T1-weighted anatomical scans with the following parameters were obtained at each TP: TR = 9.8 ms, TE = 4.6 ms, flip angel = 8°, 140 slices, 0.875 mm × 0.875 mm × 1.2 mm, and FOV = 224 × 177 × 168 mm. Scan time for this sequence was 4 min 56 s. There were no major scanner hardware or software upgrades during the MRI data collection period. A radiologist reviewed all scans at TP1 and no anomalous findings were reported.

### Image analysis

Image processing was performed on the computer network at Leiden University Medical Center. Whole-brain volumetric segmentation and cortical surface reconstruction was performed using FreeSurfer 5.3, a well-validated open-source software suite which is freely available (http://surfer.nmr.mgh.harvard.edu/). The technical details of this automated processing and the specific processing steps are described in detail elsewhere (Dale, Fischl, & Sereno, 1999; Fischl, 2012; Fischl et al., 2002; Fischl, Sereno, & Dale, 1999). Next, the images were processed using FreeSurfer 5.3’s longitudinal stream (Reuter, Schmansky, Rosas, & Fischl, 2012). Specifically, an unbiased within-subject template space and image (“base”) is created using robust, inverse consistent registration (Reuter, Rosas, & Fischl, 2010). Several processing steps, such as skull stripping, Talairach transforms, atlas registration, and spherical surface maps and parcellations are then initialized with common information from the within-subject template, significantly increasing reliability and statistical power (Reuter et al., 2012).

Detailed post-processing QC was then performed by trained operators on all scans. This QC procedure was performed prior to the hippocampal surbregion segmentation. The visual inspection focused both on overall image quality, including motion artifacts, and the accuracy of the whole-brain volumetric segmentations and the reconstructed surfaces. Scans judged to be of poor quality, either due to poor contrast or motion, or due to markedly inaccurate segmentations and/or surfaces, were excluded and the remaining scans from that participant were reprocessed through the longitudinal pipeline to assure the quality of the within-subject template. This QC procedure was repeated until only acceptable scans were included in the longitudinal processing (note that single time points were also processed longitudinally). No manual editing was performed.

Finally, using FreeSurfer 6.0, the T1-weighthed images were processed using a novel automated algorithm for longitudinal segmentation of hippocampal subregions (Iglesias et al., 2015; Iglesias et al., 2016) (Figure 1). The procedure uses a computational atlas built from high resolution *ex vivo* MRI data, acquired at an average of 0.13 mm isotropic resolution on a 7-Tesla scanner, and an *in vivo* atlas that provides information about adjacent extrahippocampal structures (Iglesias et al., 2015). The unbiased longitudinal segmentation relies on subject-specific atlases and the segmentations at the different TPs are jointly computed using a Bayesian inference algorithm (Iglesias et al., 2016). Compared with the previous algorithm developed by FreeSurfer (Van Leemput et al., 2009), the volumes generated by this new algorithm are more comparable with histologically based measurements of the subfields and much closer to the underlying subregion boundaries (Iglesias et al., 2015). It also provides a more comprehensive, fine-grained segmentation of the structures of the hippocampus. For each hemisphere, the following 12 subregions are segmented: parasubiculum, presubiculum, subiculum, CA1, CA2/3 (combined in the atlas due to indistinguishable MRI contrast), CA4, the granule cell layer of the DG (GC-DG), the molecular layer (ML), fimbria, the hippocampal fissure, the hippocampus-amygdala transition area (HATA), and the hippocampal tail (the posterior end of the hippocampus, which includes portions of the CA fields and DG undistinguishable with the MRI contrast). Test-retest reliability has been found to be high or moderate-to-high for all subregions except the hippocampal fissure in samples of older adults and young adults with T1-weigthed images with standard resolution (Whelan et al., 2016), and to be further improved for nearly all the regions by use of the longitudinal pipeline (Iglesias et al., 2016). In addition to the subregions, a measure of whole hippocampus volume is obtained by adding up the volumes of the subregions (not including the hippocampal fissure). For each scan, volumetric estimates for each annotation was extracted and averaged across hemispheres. Additionally, we extracted measures of estimated intracranial volume (ICV) from an atlas-based spatial normalization procedure (Buckner et al., 2004). Note that as FreeSurfer 5.3’s longitudinal pipeline assumes a constant ICV, the ICV measures were extracted from the cross-sectionally processed scans.

**Figure 1.**
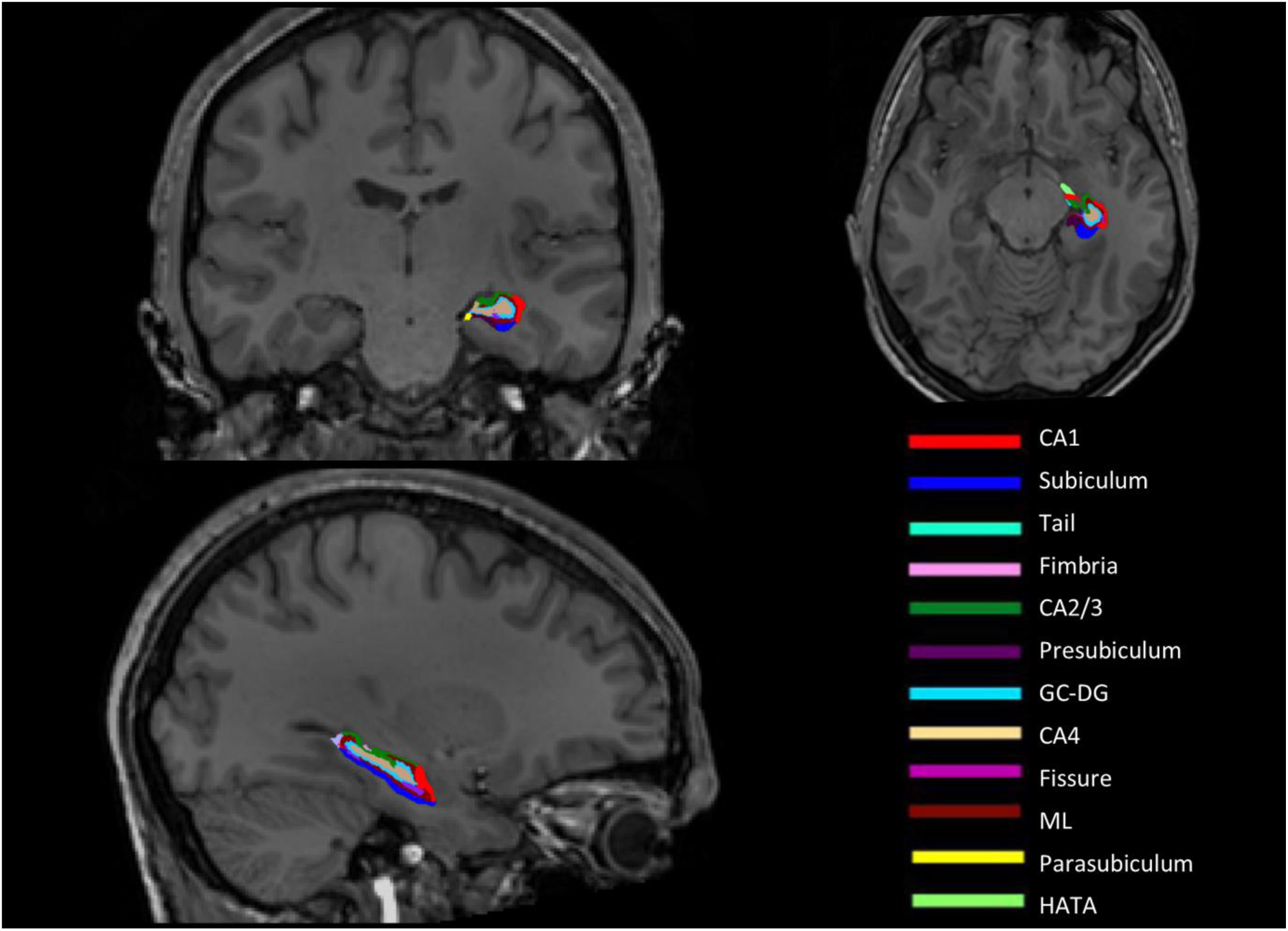
Color-coded illustration of the hippocampal subregions in coronal (top left), horizontal (top right) and sagittal (bottom left) views from a representative participant. The subregion volumes are overlaid on the whole-brain T1-weighted longitudinally processed image.

### Statistical analysis

Statistical analyses were performed using IBM SPSS 24.0 (IBM Corporation) and R 3.3.3 (https://www.r-project.org/). To test for reliability over time in our longitudinal sample, intra class correlation (ICC) was calculated for whole hippocampal volume and all subregions (Table 2). Consistent with previous reports (Whelan et al., 2016), ICC was high for all variables except the hippocampal fissure. To investigate developmental trajectories of volume of total hippocampus and each of the 12 hippocampal subregions, and the effects of sex, we used mixed models, performed using the *nlme* package (Pinheiro, Bates, DebRoy, Sarkar, & R Core Team, 2017). Mixed modelling approaches are well suited for accelerated longitudinal designs and able to handle missing data, and for these reasons widely used (Vijayakumar, Mills, Alexander-Bloch, Tamnes, & Whittle, in press). All mixed models followed a formal model-fitting procedure. Preferred models had lower Bayesian Information Criterion (BIC) values. This model selection procedure was used to ensure the most parsimonious model was selected (i.e., choosing the less complex model when the addition of parameters do not improve model fit). First, we ran an unconditional means model including a fixed and random intercept to allow for individual differences. Second, we then compared these models with three often used different growth models (linear, quadratic, and cubic (Casey, 2015)) that tested the grand mean trajectory of age using the polynomial function. Third, we added a random slope to the best fitting age model and tested whether this improved model fit. Fourth, to investigate sex differences in raw volume and volume change over time, we added sex as a main effect and an interaction effect, respectively, to the best fitting model and tested whether either of these improved model fit.

**Table 2.** Intra class correlation (ICC) for whole hippocampus and hippocampal subregion volumes

| **Region** | **ICC** |
| --- | --- |
| Whole hippocampus | 0.969 |
| Parasubiculum | 0.937 |
| Presubiculum | 0.953 |
| Subiculum | 0.962 |
| CA1 | 0.966 |
| CA2/3 | 0.948 |
| CA4 | 0.944 |
| GC-DG | 0.951 |
| ML | 0.965 |
| Fimbria | 0.867 |
| Hippocampal fissure | 0.694 |
| HATA | 0.918 |
| Hippocampal tail | 0.866 |

Notes. CA = cornu ammonis, GC-DG = granule cell layer of dentate gyrus, ML = molecular layer, HATA = hippocampus-amygdala transition area.

In a set of follow-up analyses, we added a linear growth model of ICV to the best fitting model and checked how this affected the significance for each of the age terms and sex. However, in our discussion we focus on the results for raw volumes, as we were mainly interested in how subregion volumes change over time, and how these longitudinal developmental patterns are associated with sex and general cognitive ability. First, previous results show that whether and how one includes a global variable like ICV in the statistical analyses may directly influence regional results in complex ways (Dennison et al., 2013; Pintzka, Hansen, Evensmoen, & Håberg, 2015; Sanfilipo, Benedict, Zivadinov, & Bakshi, 2004). Second, recent results also show that global metrics, including ICV, continue to change in late childhood and adolescence (Mills et al., 2016) and controlling for these measures in developmental studies thus generates a different research question of relative change. Finally, the inclusion of a global variable may be redundant when examining longitudinal change using mixed models, as each subject receives its own intercept and slope (Crone & Elzinga, 2015). Thus, the between-subject variance due to individual differences in head size is captured at the individual level over time; allowing for better characterization of changes in regional volume estimates over time (see also (Herting et al., 2018; Vijayakumar et al., in press)).

Finally, we investigated whether level of general cognitive ability could explain variance in hippocampal subregion volumes and/or development. For each participant we calculated an average general cognitive ability score across TP1 and TP2 from the T-scores on the available subtests to obtain a single score per participant. This yielded a subsample of 259 participants with 667 scans (11 participants only had MRI data included from TP3 where no IQ tasks were performed). The mean score for this sample was 109.1 (SD = 9.4, range = 80.0-147.5). We then added this continuous general cognitive ability score (centered) to the best fitting mixed model and checked the significance of its main and age interaction terms. These results were corrected for multiple comparisons using a Bonferroni procedure adjusted for correlated variables (using the mean correlation between the 13 volumes; whole hippocampal volume and the 12 hippocampal subregions) (http://www.quantitativeskills.com/sisa/calculations/bonfer.htm) (Perneger, 1998; Sankoh, Huque, & Dubey, 1997), yielding a significance level for α (2-sided adjusted) = .0144. For visualization only, the sample was split into two approximately equally large subgroups: relatively low (mean = 102.2, SD = 5.5, range = 80.0-108.8, 325 scans) and relatively high (mean = 116.1, SD = 5.5, range = 110.0-147.5, 342 scans) general cognitive ability. Finally, in follow-up analyses for subregions where the general cognitive ability main or age interaction term was significant, we reran the models after adding a linear growth term of ICV.

## Results

### Hippocampal subregion development

BIC values for the different unconditional means models and age models for the volume of the whole hippocampus and each hippocampal subregion are reported in Table 3. Model parameters for the best fitting models are reported in Table 4. Mixed model analyses on whole hippocampus volume showed a cubic developmental pattern. As shown in Figure 2, whole hippocampus volume increased in late childhood and early adolescence, followed by a slightly decelerating decrease in late adolescence and young adulthood.

**Figure 2.**
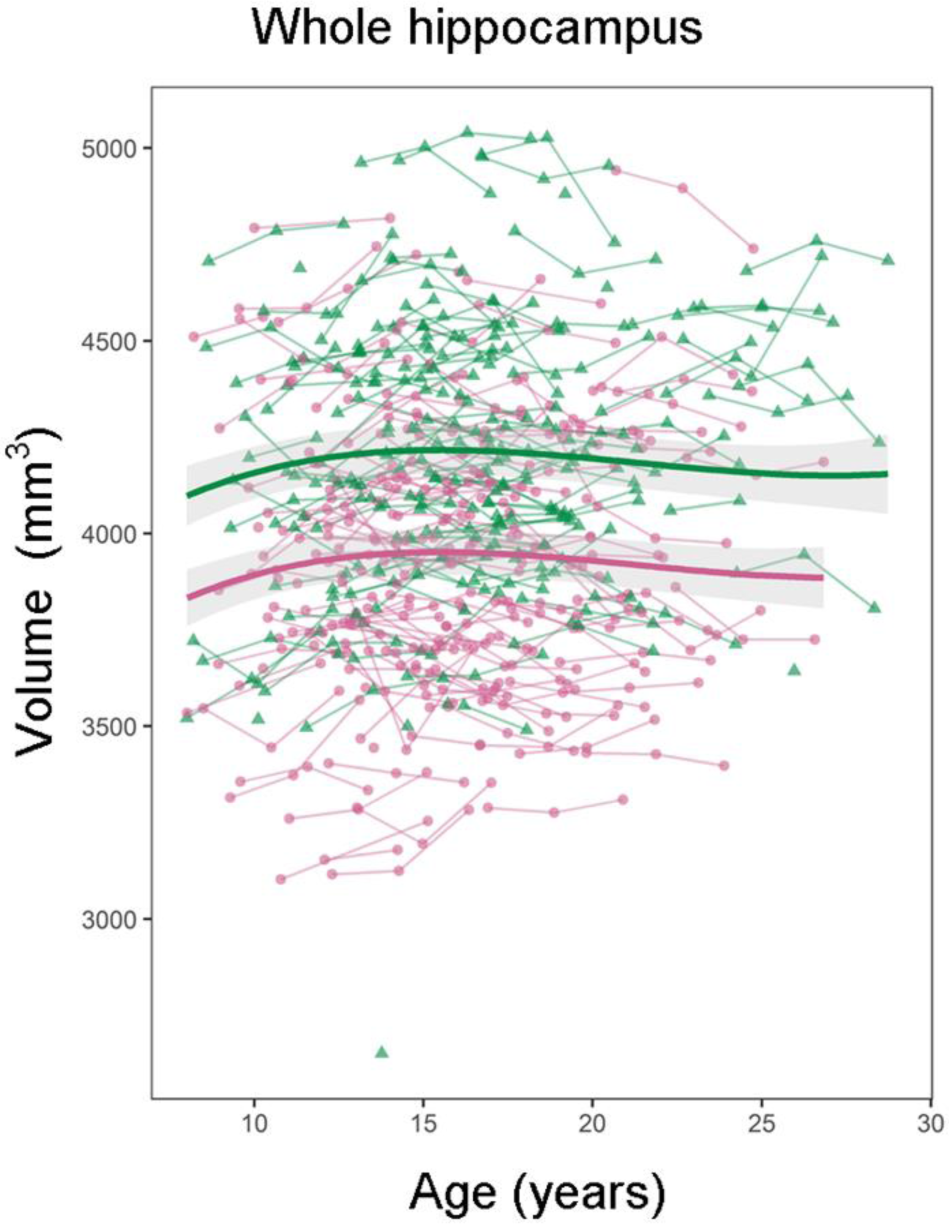
Development of whole hippocampus volume. Volume (y-axis) by age (x-axis) and the optimal fitting model, a cubic model, is shown. The shaded areas represents the 95% confidence intervals. Individual boys (green) and girls (pink) are represented by individual lines, and participants measured once are represented by dots.

**Table 3.**
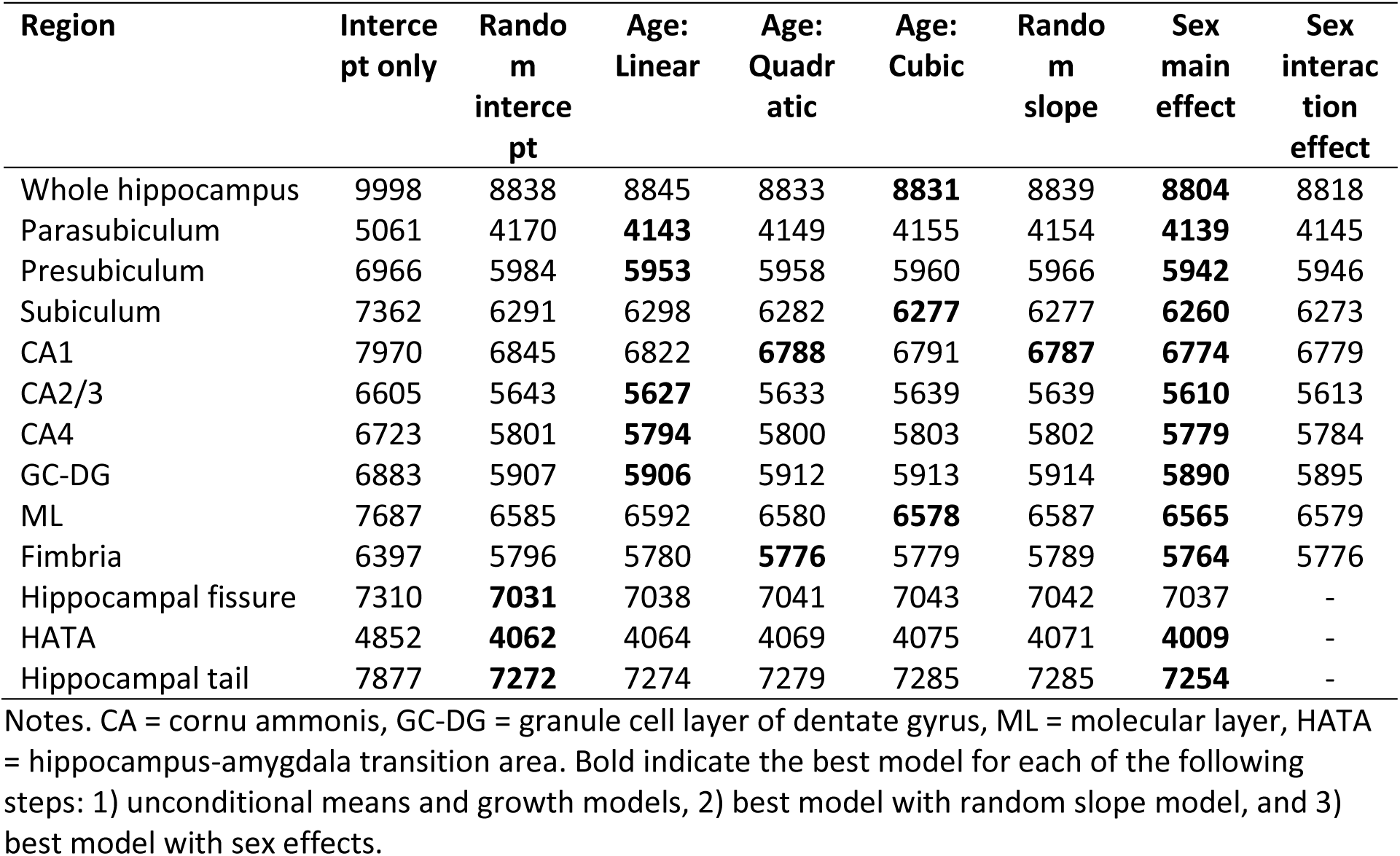
BIC values for the comparison of different mixed models examining age and sex effect on whole hippocampus and hippocampal subregion volumes

| Region | Intercept only | Random intercept | Age: Linear | Age: Quadratic | Age: Cubic | Random slope | Sex main effect | Sex interaction effect |
| --- | --- | --- | --- | --- | --- | --- | --- | --- |
| Whole hippocampus | 9998 | 8838 | 8845 | 8833 | **8831** | 8839 | **8804** | 8818 |
| Parasubiculum | 5061 | 4170 | **4143** | 4149 | 4155 | 4154 | **4139** | 4145 |
| Presubiculum | 6966 | 5984 | **5953** | 5958 | 5960 | 5966 | **5942** | 5946 |
| Subiculum | 7362 | 6291 | 6298 | 6282 | **6277** | 6277 | **6260** | 6273 |
| CA1 | 7970 | 6845 | 6822 | **6788** | 6791 | **6787** | **6774** | 6779 |
| CA2/3 | 6605 | 5643 | **5627** | 5633 | 5639 | 5639 | **5610** | 5613 |
| CA4 | 6723 | 5801 | **5794** | 5800 | 5803 | 5802 | **5779** | 5784 |
| GC-DG | 6883 | 5907 | **5906** | 5912 | 5913 | 5914 | **5890** | 5895 |
| ML | 7687 | 6585 | 6592 | 6580 | **6578** | 6587 | **6565** | 6579 |
| Fimbria | 6397 | 5796 | 5780 | **5776** | 5779 | 5789 | **5764** | 5776 |
| Hippocampal fissure | 7310 | **7031** | 7038 | 7041 | 7043 | 7042 | 7037 | - |
| HATA | 4852 | **4062** | 4064 | 4069 | 4075 | 4071 | **4009** | - |
| Hippocampal tail | 7877 | **7272** | 7274 | 7279 | 7285 | 7285 | **7254** | - |

Notes. CA = cornu ammonis, GC-DG = granule cell layer of dentate gyrus, ML = molecular layer, HATA = hippocampus-amygdala transition area. Bold indicate the best model for each of the following steps: 1) unconditional means and growth models, 2) best model with random slope model, and 3) best model with sex effects.

**Table 4.**
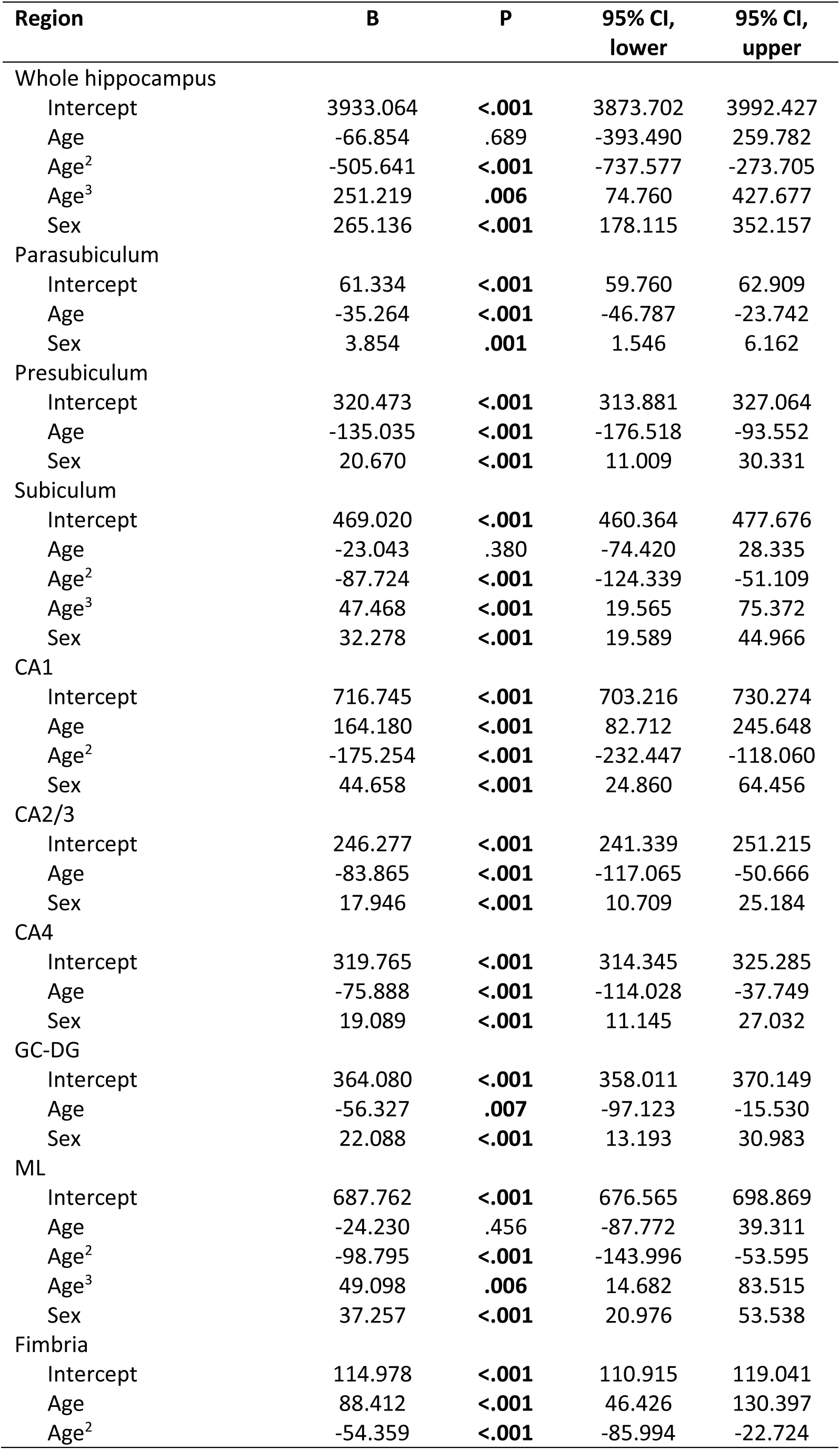
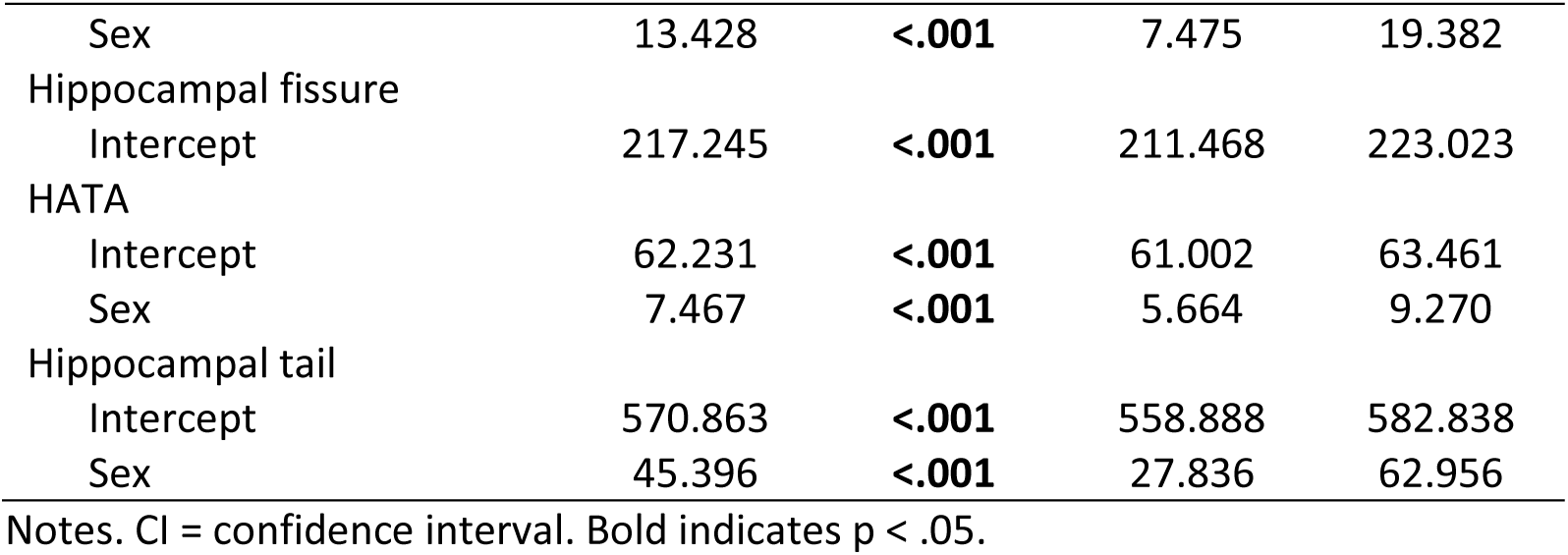
Model parameters for fixed effects in the best fitting model for whole hippocampus and hippocampal subregion volumes

Best fitting models for all hippocampal subregions are shown in Figure 3. For parasubiculum, presubiculum, CA2/3, CA4 and GC-DG, a linear age model fitted best, with steady volume decreases from late childhood to adulthood. For CA1, a quadratic age model with random slope fitted best, and the quadratic age model was also the best fit for fimbria. For both of these subregion volumes, development followed an inverse-u trajectory. For subiculum and ML volumes, development followed a cubic pattern similar to whole hippocampus volume; early increases, followed by decelerating decreases. Finally, for the three subregions the hippocampal fissure, HATA and the hippocampal tail, the random intercept model fitted better than any of the growth models.

**Figure 3.**
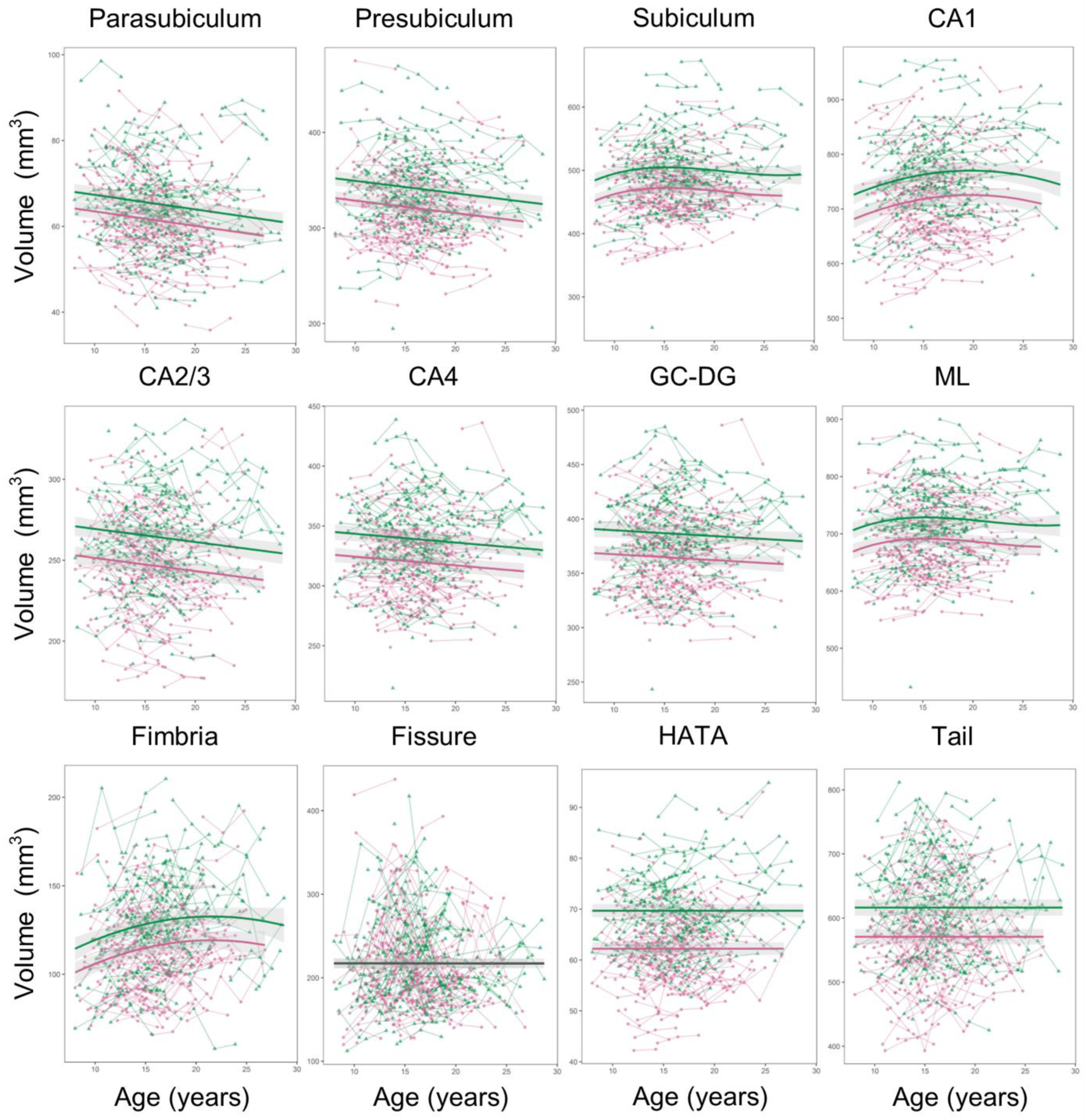
Development of hippocampal subregions. Volumes (y-axis) by age (x-axis) and the optimal fitting models are shown. A linear model fitted best for parasubiculum, presubiculum, CA2/3, CA4 and GC-DG, a quadratic model fitted best for CA1 and fimbria, a cubic model fitted best for subiculum and ML, and a random intercept model fitted best for the hippocampal fissure, HATA and the hippocampal tail. There was a main effect of sex for all subregions except the hippocampal fissure, but no interaction effects between sex and age. The shaded areas represents the 95% confidence intervals. Individual boys (green) and girls (pink) are represented by individual lines, and participants measured once are represented by dots.

### Sex effects on hippocampal subregion volumes and development

Both for whole hippocampus volume and for all subregions except the hippocampal fissure, adding sex as a main effect improved model fit (Table 3). In all these regions, boys on average had larger volume than girls (Table 4, Figures 2 and 3). However, adding sex as an interaction effect did not improve model fit for whole hippocampus volume or any of the hippocampal subregions. This indicates parallel developmental trajectories in girls and boys.

### ICV adjusted results

In order to better be able to compare our results with some of the previous studies, we added a linear growth model of ICV to the best fitting model for whole hippocampus and each subregion volume (Table 5). ICV was significant for all regions except the hippocampal fissure, while the effect of sex was no longer significant in any region except for whole hippocampus and HATA. For the subregions, most of the age effects remained significant, with the exception of the linear age term for GC-DG and the cubic age term for subiculum and ML.

**Table 5.** Model parameters for fixed effects when including a linear growth model of ICV in the best fitting models for whole hippocampus and hippocampal subregion volumes

| **Region** | **B** | **P** |
| --- | --- | --- |
| Whole hippocampus |  |  |
| Intercept | 4002.626 | **<.001** |
| Age | 289.818 | .093 |
| Age<sup>2</sup> | -227.507 | .068 |
| Age<sup>3</sup> | 123.706 | .185 |
| Sex | 115.089 | **.006** |
| ICV | 3256.565 | **<.001** |
| Parasubiculum |  |  |
| Intercept | 62.147 | **<.001** |
| Age | -32.475 | **<.001** |
| Sex | 2.090 | .099 |
| ICV | 38.415 | **.001** |
| Presubiculum |  |  |
| Intercept | 326.265 | **<.001** |
| Age | -112.343 | **<.001** |
| Sex | 8.124 | .107 |
| ICV | 272.919 | **<.001** |
| Subiculum |  |  |
| Intercept | 479.746 | **<.001** |
| Age | 32.442 | .222 |
| Age<sup>2</sup> | -45.678 | **.018** |
| Age<sup>3</sup> | 27.947 | .053 |
| Sex | 9.111 | .146 |
| ICV | 502.530 | **<.001** |
| CA1 |  |  |
| Intercept | 729.120 | **<.001** |
| Age | 226.468 | **<.001** |
| Age<sup>2</sup> | -120.395 | **<.001** |
| Sex | 17.975 | .067 |
| ICV | 587.351 | **<.001** |
| CA2/3 |  |  |
| Intercept | 251.240 | **<.001** |
| Age | -62.749 | **<.001** |
| Sex | 7.227 | .052 |
| ICV | 232.916 | **<.001** |
| CA4 |  |  |
| Intercept | 326.083 | **<.001** |
| Age | -50.078 | **.011** |
| Sex | 5.483 | .172 |
| ICV | 295.554 | **<.001** |
| GC-DG |  |  |
| Intercept | 370.766 | **<.001** |
| Age | -30.442 | .149 |
| Sex | 7.663 | .089 |
| ICV | 313.704 | **<.001** |
| ML |  |  |
| Intercept | 699.541 | **<.001** |
| Age | 38.222 | .256 |
| Age<sup>2</sup> | -51.769 | **.034** |
| Age<sup>3</sup> | 27.628 | .130 |
| Sex | 11.898 | .138 |
| ICV | 550.079 | **<.001** |
| Fimbria |  |  |
| Intercept | 118.896 | **<.001** |
| Age | 99.889 | **<.001** |
| Age<sup>2</sup> | -41.481 | **.011** |
| Sex | 4.895 | .147 |
| ICV | 185.290 | **<.001** |
| Hippocampal fissure |  |  |
| Intercept | 217.206 | **<.001** |
| ICV | 107.204 | .131 |
| HATA |  |  |
| Intercept | 63.359 | **<.001** |
| Sex | 4.994 | **<.001** |
| ICV | 54.358 | **<.001** |
| Hippocampal tail |  |  |
| Intercept | 581.907 | **<.001** |
| Sex | 21.282 | **.033** |
| ICV | 529.434 | **<.001** |

Notes. Bold indicates $p < .05$ .

### General cognitive ability and hippocampal subregion volumes and development

To investigate whether level of general cognitive ability could explain variance in hippocampal volumes and/or development, we added this continuous score as an interaction term to the best fitting model (Table 6). Significant positive main effects of general cognitive ability were found for two subregions: CA2/3 (B = 0.568, p = .004) and CA4 (B = 0.695, p =.001), such that higher level of performance was related to great volumes. Additionally, there was an uncorrected positive effect for GC-DG (B = 0.513, p =.037). The results also revealed a significant quadratic age × general cognitive ability interaction for ML (B = −6.937, p =.012), and a similar uncorrected effect for subiculum (B = - 4.569, p =.042). The results of these analyses with general cognitive ability as a continuous measure are illustrated using subgroups of relatively low and relatively high general cognitive ability (Figure 4). For these subregions where the general cognitive ability main or an age interaction term was significant, we reran the models with a linear growth term of ICV. The positive main effects of general cognitive ability remained significant for CA2/3 (B = 0.459, p = .012) and CA4 (B = 0.552, p =.005), while the previously uncorrected positive effect for GC-DG was no longer significant (B = 0.360, p =.109). After adding a linear trend of ICV, the quadratic age × general cognitive ability interaction showed an uncorrected effect for ML (B = −6.195, p =.028), and was no longer significant for subiculum (B = −3.821, p =.087).

**Figure 4.**
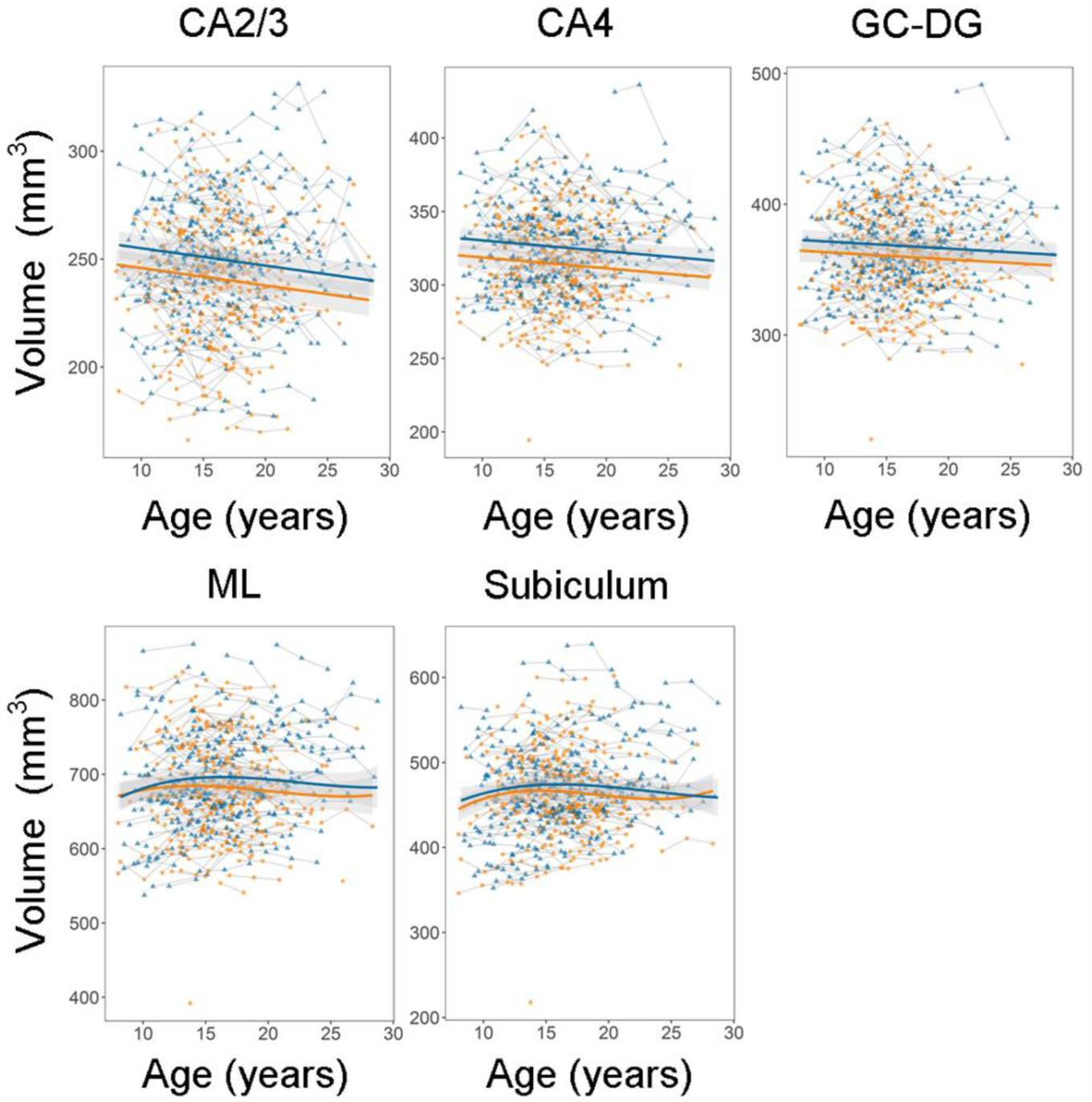
Associations between general cognitive ability and hippocampal subregion volumes and development. For visualization purposes, the sample was split into two groups: relatively high (blue) and relatively low (orange) general cognitive ability. Note that the statistical analyses were performed using a continuous general cognitive ability score, and by adding this score as an interaction term to the best fitting model. Volumes (y-axis) by age (x-axis) are shown and the shaded areas represents the 95% confidence intervals.

**Table 6.** Model parameters for fixed effects when including level of general cognitive ability in the best fitting models for whole hippocampus and hippocampal subregion volumes

| **Region** | **B** | **P** |
| --- | --- | --- |
| **Whole hippocampus** |  |  |
| Intercept | 3924.655 | **<.001** |
| Age | -3.352 | .984 |
| Age<sup>2</sup> | -424.260 | **<.001** |
| Age<sup>3</sup> | 302.973 | **.002** |
| Sex | 278.766 | **<.001** |
| GCA | 4.165 | .082 |
| Age × GCA | -5.571 | .788 |
| Age<sup>2</sup> × GCA | -25.242 | .077 |
| Age<sup>3</sup> × GCA | -13.688 | .198 |
| **Parasubiculum** |  |  |
| Intercept | 61.129 | **<.001** |
| Age | -35.298 | **<.001** |
| Sex | 3.915 | **.001** |
| GCA | 0.013 | .835 |
| Age × GCA | 0.250 | .711 |
| **Presubiculum** |  |  |
| Intercept | 319.470 | **<.001** |
| Age | -135.823 | **<.001** |
| Sex | 20.163 | **<.001** |
| GCA | -0.057 | .827 |
| Age × GCA | 3.009 | .214 |
| **Subiculum** |  |  |
| Intercept | 467.638 | **<.001** |
| Age | -11.672 | .662 |
| Age<sup>2</sup> | -72.576 | **<.001** |
| Age<sup>3</sup> | 54.873 | **<.001** |
| Sex | 33.274 | **<.001** |
| GCA | 0.500 | .154 |
| Age × GCA | 1.539 | .635 |
| Age<sup>2</sup> × GCA | -4.569 | **.042** |
| Age<sup>3</sup> × GCA | -2.734 | .103 |
| **CA1** |  |  |
| Intercept | 713.034 | **<.001** |
| Age | 163.514 | **<.001** |
| Age<sup>2</sup> | -162.828 | **<.001** |
| Sex | 49.062 | **<.001** |
| GCA | 0.830 | .122 |
| Age × GCA | 7.095 | .158 |
| Age<sup>2</sup> × GCA | -4.528 | .189 |
| **CA2/3** |  |  |
| Intercept | 245.720 | **<.001** |
| Age | -83.888 | **<.001** |
| Sex | 19.013 | **<.001** |
| GCA | 0.568 | **.004** |
| Age × GCA | 0.967 | .618 |
| **CA4** |  |  |
| Intercept | 320.313 | **<.001** |
| Age | -76.403 | **<.001** |
| Sex | 19.165 | **<.001** |
| GCA | 0.695 | **.001** |
| Age × GCA | 0.602 | .787 |
| GC-DG |  |  |
| Intercept | 364.380 | **<.001** |
| Age | -57.441 | **.006** |
| Sex | 22.351 | **<.001** |
| GCA | 0.513 | **.037** |
| Age × GCA | 0.542 | .820 |
| ML |  |  |
| Intercept | 687.289 | **<.001** |
| Age | -12.671 | .701 |
| Age<sup>2</sup> | -78.680 | **.001** |
| Age<sup>3</sup> | 56.778 | **.003** |
| Sex | 39.175 | **<.001** |
| GCA | 0.754 | .094 |
| Age × GCA | 2.038 | .611 |
| Age<sup>2</sup> × GCA | -6.937 | **.012** |
| Age<sup>3</sup> × GCA | -2.543 | .218 |
| Fimbria |  |  |
| Intercept | 114.412 | **<.001** |
| Age | 86.443 | **<.001** |
| Age<sup>2</sup> | -50.670 | **.003** |
| Sex | 14.347 | **<.001** |
| GCA | -0.062 | .711 |
| Age × GCA | 3.458 | .176 |
| Age<sup>2</sup> × GCA | -1.378 | .467 |
| Hippocampal fissure |  |  |
| Intercept | 217.184 | **<.001** |
| GCA | -0.599 | .066 |
| HATA |  |  |
| Intercept | 61.972 | **<.001** |
| Sex | 7.791 | **<.001** |
| GCA | 0.002 | .960 |
| Hippocampal tail |  |  |
| Intercept | 569.507 | **<.001** |
| Sex | 49.356 | **<.001** |
| GCA | 0.551 | .256 |

Notes. GCA = general cognitive ability. Bold indicates $p < .05$ .

## Discussion

The current study of longitudinal development of hippocampal subregions from childhood to adulthood yielded three novel findings. First, the results showed heterogeneous developmental patterns across subregions, with nonlinear trajectories with early volume increases for subiculum, CA1, ML and fimbria, and linear volume decreases or no change in the other subregions. Second, boys showed larger volumes than girls for almost all hippocampal subregions, but boys and girls showed parallel developmental trajectories. Third, general cognitive ability was positively associated with CA2/3 and CA4 volumes and with ML development. These findings will be discussed in more detail in the following paragraphs.

Whole hippocampal volume increased in late childhood and early adolescence, followed by a slightly decelerating decrease in late adolescence and young adulthood, in agreement with accumulating evidence from other studies (Coupe et al., 2017; Herting et al., 2018; Narvacan et al., 2017; Wierenga et al., 2014). Most importantly, however, distinct hippocampal subfields showed different developmental trajectories. Subiculum, CA1, ML and fimbria showed nonlinear trajectories with initial volume increases. In stark contrast, parasubiculum, presubiculum, CA2/3, CA4 and GC-DG showed linear volume decreases. Finally, the hippocampal fissure, HATA and the hippocampal tail showed no development across adolescence.

Our results appear to be consistent with the observed age-related increase in CA1 in the right hemisphere in late childhood and early adolescence in the study by Lee et al., but not with the observed age-related increase in the right CA3/DG in the same study (Lee, Ekstrom, et al., 2014). Compared to the results by Daugherty et al., our results appear consistent with the observed negative age relationship for CA3/DG volume, but partly at odds with the observed negative age relationship for CA1/2 volume (Daugherty et al., 2016). Direct comparisons between our developmental results and previous studies of specific hippocampal subregions (Daugherty et al., 2016; Krogsrud et al., 2014; Lee, Ekstrom, et al., 2014; Tamnes et al., 2014) are however difficult, as the previous studies relied on small and/or cross-sectional samples of children and adolescents.

Additionally, two of the previous studies (Daugherty et al., 2016; Lee, Ekstrom, et al., 2014) relied on manual segmentation with its limitations; being laborious and liable to bias and variability (Schlichting, Mack, et al., 2017). Two other previous studies (Krogsrud et al., 2014; Tamnes et al., 2014) used an older automated segmentation procedure which has been found to systemically misestimate specific subregion volumes compared to histological classifications (Schoene-Bake et al., 2014) and for many subregions to show poor agreement with the newer automated procedure used in the present study (Whelan et al., 2016).

We were also interested in testing sex differences in trajectories of hippocampal development. Boys showed larger volumes than girls for all hippocampal subregions except the hippocampal fissure, but adding sex as an interaction term did not improve model fit for any region. Our results therefore do not indicate sex differences in the development of hippocampal subregion volumes. Early cross-sectional studies of whole hippocampal volume reported conflicting sex-specific age-related differences (Giedd et al., 1996; Suzuki et al., 2005), but larger or longitudinal studies have not found sex differences in developmental trajectories (Dennison et al., 2013; Koolschijn & Crone, 2013; Wierenga et al., 2014). The present results are also consistent with the previous studies on hippocampal subregions which have found larger absolute volumes in boys (Krogsrud et al., 2014; Tamnes et al., 2014), but no interactions between sex and age (Krogsrud et al., 2014) or sex differences in change rates (Tamnes et al., 2014). Notably, and consistent with several previous reports on sex differences in brain volumes (Marwha, Halari, & Eliot, 2017; Pintzka et al., 2015; Tan, Ma, Vira, Marwha, & Eliot, 2016) and as expected, most of the main effects of sex on hippocampal subregion volumes disappeared when including ICV in the statistical models, indicating that sex plays a minor role for hippocampal subregion volume differences. Studies investigating effects of puberty and sex hormones on hippocampal subregion development are however needed (see (Herting & Sowell, 2017) and discussion of future directions below).

Functionally, it is likely that different parts of the hippocampus have somewhat different roles for different aspects of cognition and behavior. Our results showed that higher general cognitive ability was associated with greater CA2/3 and CA4 volumes across the investigated age-span. Additionally, general cognitive ability was also associated with the developmental trajectory for ML volume, such that individuals with higher scores showed a slightly more nonlinear development. A similar association has previously been found between general intellectual ability and cortical development (Shaw et al., 2006). Previous studies of hippocampal subregion volumes and development in children and adolescents have focused on associations with learning and memory (Daugherty, Flinn, & Ofen, 2017; DeMaster et al., 2014; Lee, Ekstrom, et al., 2014; Riggins, Blankenship, Mulligan, Rice, & Redcay, 2015; Schlichting, Guarino, Schapiro, Turk-Browne, & Preston, 2017; Tamnes et al., 2014). For instance, a recent study found that a multivariate profile of age-related differences in intrahippocampal volumes was associated with differences in encoding of unique memory representations (Keresztes et al., 2017). The hippocampus does however appear to be involved in a broad specter of cognitive functions and behaviors that may also include e.g. spatial navigation, emotional behavior, stress regulation, imagination and prediction (Aribisala et al., 2014; Lee, Johnson, & Ghetti, 2017; Mullally & Maguire, 2014; Rubin, Watson, Duff, & Cohen, 2014). Intriguingly, in a large study of older adults, general intelligence was found to be associated with several measures of tissue microstructure in the hippocampus, which were derived from diffusion tensor imaging, magnetization transfer and relaxometry, but not with whole hippocampus volume (Aribisala et al., 2014). This suggested that more subtle differences in the hippocampus may reflect differences in general cognitive ability, at least in the elderly (see also (Reuben, Brickman, Muraskin, Steffener, & Stern, 2011)). Our results add to this picture by indicating that specific hippocampal subregion volumes and developmental patterns may be associated with general cognitive ability in youth.

Our study has several strengths, including a large sample size, a longitudinal design with up to three scans per participant, the use of a new hippocampal subregion segmentation tool, and longitudinal image processing; however, there are also important limitations that need to be considered. An urgent limitation is that we used only T1-weighthed data acquired on a 3-Tesla scanner with standard resolution (0.875×0.875×1.2 mm). Strongly preferable, scans with higher spatial resolution should be used, and it is also better to use a combination of T1-weighted and T2-weighted data to improve contrast (Iglesias et al., 2015). The method employed to segment hippocampal subregions was developed based on *ex vivo* tissues scanned with ultra-high field strength, and has been demonstrated to be applicable and reliable in datasets with different types of resolution and contrast (Iglesias et al., 2015; Whelan et al., 2016). Nonetheless, our results, particularly for the hippocampal fissure which showed relatively lower reliability over time and for the smaller subregions (e.g., parsubiculum, HATA and fimbria), should be interpreted with caution. Future longitudinal developmental studies with higher resolution scans are called for. A second caveat is that there is disagreement across both manual and automated segmentation methods about the placement of certain subregion boundaries (Wisse et al., 2017; Yushkevich, Amaral, et al., 2015). Direct comparisons between the new FreeSurfer automated method used in the present study and other available automated methods such as Automatic Segmentation of Hippocampal Subfields (ASHS) (Yushkevich, Pluta, et al., 2015), Multiple Automatically Generated Templates (MAGeT) (Pipitone et al., 2014) and Advanced Neuroimaging Tools (ANTs) (Avants et al., 2011), are also lacking (see (Schlichting, Mack, et al., 2017) for such a study, comparing ASHS, ANTs and manual segmentation in child, adolescent, and adult age groups), but critical in order to test reproducibility of the present results across available tools. Importantly, a current international collaborative effort, The Hippocampal Subfield Group (http://www.hippocampalsubfields.com/), is underway to develop a harmonized segmentation protocol to overcome this barrier (Wisse et al., 2017). Third, the hippocampal subregion segmentation method employed has not been specifically developed or validated for children or adolescents, and might be biased towards brains of older adults. Fourth, we did not investigate longitudinal change in general cognitive ability or more specific cognitive functions. Future studies are needed to further shed light on the functional implications of longitudinal changes in hippocampal subregions, both in terms of development of cognitive functions, and the emergence of mental disorder such as psychosis and depression during adolescence. Finally, we also note that our conclusions from group-level inferences may not translate to individual development, and that appropriate disambiguation of between- and within-person effects in analyses is an issue that deserves more attention in the developmental cognitive neuroscience field (Foulkes & Blakemore, 2018).

Future studies should investigate puberty and sex hormone effects on development of hippocampal subregion volumes, as it has been found that age and pubertal development have both independent and interactive influences on hippocampus volume change over adolescence (Goddings et al., 2014; Satterthwaite et al., 2014), and that puberty related increases in testosterone level are related to development of hippocampus volume in both males and females (Wierenga et al., 2018) (but see (Herting et al., 2014)). Further, a recent study showed greater variance in males than females for several brain volumes including the hippocampus (Wierenga et al., in press), and future studies should investigate whether such variability differences are general or specific for distinct hippocampal subregion volumes and development. Next, future developmental studies should integrate subregion segmentation in the transverse plane and along the longitudinal axis of the hippocampus (Lee et al., 2017). Finally, future studies could also investigate development of hippocampal-cortical networks at the level of specific hippocampal subregions, e.g. by analyzing structural covariance (Walhovd et al., 2015), structural connectivity inferred from diffusion MRI (Wendelken et al., 2015), or functional connectivity from functional MRI (Blankenship, Redcay, Dougherty, & Riggins, 2017; Paz-Alonso, Gallego, & Ghetti, 2013).

In conclusion, our results indicate that hippocampal subregions develop in diversified ways across adolescence, with nonlinear trajectories with early volume increases for subiculum, CA1, ML and fimbria, and linear volume decreases for parasubiculum, presubiculum, CA2/3, CA4 and GC-DG. Further, while boys had larger hippocampal subregion volumes than girls, we found no sex differences in the development of the subregions. The results also indicate that volume and developmental pattern of specific hippocampal subregions may be associated with general cognitive ability. However, future studies validating the use of the employed hippocampal subregion segmentation method in samples of youth and studies directly comparing this method with other automated segmentation methods are needed, and, critically, longitudinal developmental studies with high-resolution scans are called for.

## Acknowledgements

This study was supported by the Research Council of Norway and the University of Oslo (FRIMEDBIO 230345 to CKT), and the European Research Council Starting Grant scheme (ERC-2010-StG_263234 to EAC).

